# Management of Root-Knot Nematode-Fusarium Wilt Complex with different Bio-management methods in Tomato (*Solanum lycopersicum L.*) var Srijana

**DOI:** 10.64898/2026.10.25.761656

**Authors:** Anup Bhatt, Aditi Bhandari, Sajan Gaudel, Shalik Ram Adhikari, Suman Dhakal, Ankita Baral, Sparsha Naharki, Priya Shrestha, Deepa Koirala

**Author notes:** **Corresponding author:**Anup Bhatt, Faculty of Agriculture, Agriculture and Forestry University, Chitwan, Nepal, **Email:**. Authors’ details.

## Abstract

The root-knot nematode-*Fusarium* complex is a major constraint to protected tomato cultivation in Nepal, particularly on the widely grown Srijana hybrid, which lacks Mi-mediated nematode resistance. The present study was conducted to identify the most effective biocontrol agent, to identify the most effective *Solanum* rootstock, and to compare the overall efficacy of grafting versus biocontrol agents under polyhouse conditions. For the management of the root knot nematode-Fusarium complex, a pot experiment was set up in a completely randomized design at Pokhara, Kaski. Treatments comprised *Trichoderma viride*, *Pseudomonas fluorescens*, their half-dose combination, self-grafted Srijana, and four resistant rootstocks (*S. sisymbriifolium*, *S. melongena*, *S. torvum,* and VNRRs-10), together with nematode-only, *Fusarium*-only, complex (Nematode + *Fusarium*), and disease-free controls. Each plant was inoculated with approximately 2,000 nematode eggs, followed by 25 ml of 10□ spores mL□¹ of *Fusarium* seven days later to mimic the field complex. Plant growth parameters and nematode infestation data were recorded and analyzed via RStudio v. 2026.04.0. The root knot nematode-*Fusarium* complex caused significantly greater reductions in vegetative biomass than root knot nematode and *Fusarium* alone (20.32 gm vs 38.40 gm vs 51.32 gm) at 45 DAT. Single-agent biocontrol gave intermediate protection (*T. viride* 64.9%; *P. fluorescens* 47.9% nematode management efficiency), whereas combined *T. viride* + *P. fluorescens* treatment produced the highest vegetative biomass but the weakest nematode suppression (320 galls, reproduction factor: 38). Grafting onto *S. torvum* and *S. melongena* resulted in the greatest nematode suppression, achieving 97.9% and 93.6% management efficiency, respectively, while *S. torvum* alongside VNRRs-10 maintained the highest *Fusarium* suppression (92.5%). Overall resistant rootstocks consistently outperformed biological control in reducing disease severity. These findings indicate that grafting onto resistant solanum rootstocks, particularly *S. torvum* and *S. melongena*, was observed as the most effective strategy for managing the root knot nematode-*Fusarium* complex under protected cultivation, offering a viable chemical-free management option.

## Introduction

Tomato (*Solanum lycopersicum* L.) is among the most economically and nutritionally important vegetable crops worldwide (Hobson & Grierson, 1993; Kallo, 1993), and in Nepal, it ranks as the third most important vegetable crop, after cauliflower and cabbage (MoAFE, 2026). The hybrid cultivar ‘Srijana’, released through the Nepal Agriculture Research Council (NARC), dominates commercial production in Nepal because of its adaptability to plastic tunnel cultivation and tolerance to bacterial wilt (Chapagain et al., 2011; Magar et al., 2016). However, tunnel structure also elevates soil temperature (Hamal et al., 2025; Frey et al., 2020a), which creates favourable conditions for root-knot nematode (Frey et al., 2020a) and aligns with known temperature and moisture thresholds that favour Fusarium wilt (McGovern, 2015).

Root-knot nematodes (*Meloidogyne spp.*) are among the most economically damaging plant-parasitic nematode genera, with global crop losses estimated at USD 175-200 billion annually (Khan, 2023). In tomato, infection induces root galling and giant-cell formation that disrupt water and nutrient uptake (El-Sappah et al., 2019), with yield losses ranging from 25% to complete crop failure depending on cultivar susceptibility and nematode pressure (Seid et al., 2015); in Nepal, RKN-associated losses of 30-40% have been documented (Baidya et al., 2017; Rawal, 2020). *Fusarium oxysporum* fsp *lycopersici* independently causes comparable damage, with 30-40% yield loss (up to 80% under favourable conditions) and 22-55% field incidence reported across South Asian production systems (Nirmaladevi et al., 2016). Neither pathogen typically acts alone: RKN-induced root damage predisposes plants to secondary Fusarium infection, and the resulting root-knot nematode-*Fusarium* wilt disease causes earlier onset and greater yield loss than either pathogen individually (Meena et al., 2021; Yaseen et al., 2024).

Management options for this disease complex remain limited due to constraints in both host resistance and disease management strategies. Long-term application of nematicides and fungicides carries environmental and health costs, while cultivars having *Mi-1* resistance genes used against RKN are temperature-sensitive above 28°C (Wubie & Temesgen, 2019), which is precisely the temperature range that exists under polyhouse conditions. The persistence of Fusarium’s soil-borne chlamydospore further limits crop rotation as a viable control measure (Rahman et al., 2021). These constraints have shifted attention towards resistant rootstock grafting and biocontrol agents as more durable alternatives (Wang & Li, 2019).

Srijana itself, despite its dominance among tunnel growers (Nakarmi et al., 2025), lacks nematode resistance ability (Baidya et al., 2017; KC et al., 2021), a documented gap that leaves the most widely grown cultivar susceptible to both components of the disease complex. Existing Nepal-based work has only evaluated RKN management options individually (Baidya et al., 2017; Nakarmi et al., 2025; Thakur et al., 2024), but none has evaluated it under combined root-knot nematode-*Fusarium* infection.

This study addresses that gap by evaluating resistant rootstocks and biocontrol agents individually and in combination against the disease complex under polyhouse conditions, identifying the most effective option at each disease severity level and characterizing their relative performance under combined infection.

## Materials and methods

### Greenhouse test

The experiment was conducted under semi-controlled conditions (polyhouse) at the Agribusiness Promotion Training and Resource Centre (APTRC), Malepatan-5, Pokhara Metropolitan, Kaski, Nepal, at approximately 28.2191° N latitude and 83.9684° E longitude. Grafted tomato seedlings were obtained from Graft Nepal Pvt. Ltd., while non-grafted seedlings were grown on-site. The seedling growing medium was prepared by combining soil, peat moss, vermicompost, and sand in equal parts (1:1:1:1). Seed seeds were sown in the medium placed into plug germination trays and watered once a day to keep the medium adequately moist until the seedlings reached transplantable size. The experiment was set up in a polyhouse, using growing bags (10 × 20 cm) as containers. The bags were sterilized with 1% sodium hypochlorite before being filled with soil. The soil was sieved, moistened, and treated uniformly with formaldehyde (10 L/m²) using a knapsack sprayer, then heaped and covered with polythene for 3 days to retain fumigant vapor. After fumigation, the soil was turned and left exposed for 10–15 days until no detectable formalin odor remained. After that, 10 kg of sterilized soil was filled into each growing bag for seedling raising. Tomato seedlings were inoculated with root-knot nematode one week after transplanting, followed by *Fusarium* inoculation a week later (Khan & Sharma, 2020; Göze Özdemir *et al*., 2022). The trial comprised three replications, each containing three sample plants per pot. Roots were uprooted and examined at 45, 60, and 75 days after transplanting (DAT).

### Experimental design and treatment

The experiment consisted of 12 treatments arranged in a completely randomized design with three replications. Treatments T1–T3 consisted of biocontrol agents applied in combination with root-knot nematode and *Fusarium oxysporum*. T1 received *Trichoderma viride*, T2 received *Pseudomonas fluorescens*, and T3 received a combination of *T. viride* and *P. fluorescens*, each under combined nematode–*Fusarium* inoculation. T4 consisted of self-grafted Srijana under combined nematode–*Fusarium* inoculation, whereas T5–T8 consisted of Srijana scion grafted onto *Solanum sisymbriifolium*, *Solanum melongena* (Solmel), *Solanum torvum*, and VNRRs-10 rootstocks, respectively, followed by combined nematode–*Fusarium* inoculation. T9, T10, and T11 represented nematode-only, *Fusarium*-only, and combined nematode + *Fusarium* inoculated controls, respectively, while T12 was maintained as a disease-free control.

Tiny holes were made around the root zone of each plant, and roughly 2000 eggs of *Meloidogyne* spp. were introduced into the soil near the root system (Silva et al., 2017). For the inoculation of *Fusarium*, a spore suspension was prepared from actively growing PDA cultures and standardized to 10□ spores mL□¹ using a Neubauer improved hemocytometer; about 25 mL of this suspension was then applied close to each plant’s root zone to achieve even pathogen distribution across the rhizosphere (Hernandez-Aparicio et al., 2021).

Plant growth was assessed by measuring plant height (base to shoot tip) and fresh weights of root, shoot, and leaves using an electronic balance immediately after harvest, with vegetative weight calculated as their sum. Nematode infestation was evaluated through the number of egg masses per root system (the fresh weight of each root sample was weighed). Samples weighing 10 gm or less were treated with 200 mL of 0.1% sodium hypochlorite (NaOCl) to facilitate extraction of eggs from the root tissue. The resulting suspension was passed through a sieve to remove root debris, then filtered to obtain a clean egg suspension. 1 mL of this suspension was pipetted onto gridded Petri plates, which served as counting slides for microscopic examination of the eggs. This counting procedure was repeated five times per sample, and the average of these five counts was taken as the representative egg density per mL. Eggs per gram of root tissue were then determined using the unitary method:

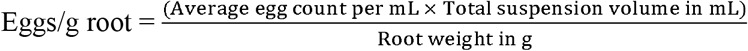

Egg count per gram of root (via extraction and microscopic counting), and the Reproduction Factor (RF), calculated following Oostenbrink (1966) as

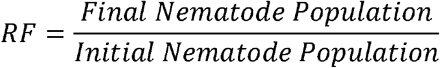

Root-knot nematode severity was scored using the 0–10 gall index of Bridge and Page (1980), while Fusarium wilt severity was scored using the 0–5 disease index of Lebeda and Buczkowski (1986). From these ratings, Disease Severity Index (DSI %) was calculated following Wheeler (1969) as:

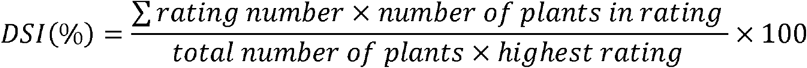

To capture cumulative disease pressure across the trial rather than severity at a single time point, the Area Under the Disease Progress Curve (AUDPC) was calculated by the trapezoidal method (Campbell & Madden, 1990) using gall index scores recorded at 45, 60, and 75 DAT, and Fusarium wilt index scores recorded at 15, 30, 45, and 60 DAI:

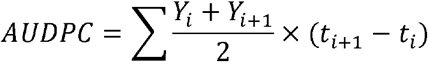

Where Yi and Y_i+1_ denote the disease index value recorded at two consecutive assessment dates, t_i_ and t_i+1_, respectively, and the summation is taken across all the successive intervals. Replicate-level AUDPC values were subjected to ANOVA, with treatment means separated by DMRT (p = 0.05). Management efficiency of each treatment was then derived from AUDPC values relative to the untreated inoculated check, following Abbott’s (1925) formula:

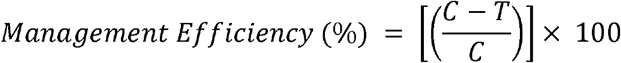

where *C* = AUDPC of the untreated inoculated check and *T* = AUDPC of the treatment.

### Pathogenicity test of *Meloidogyne* spp.

A pathogenicity test was conducted to confirm the ability of the isolated *Meloidogyne* spp. to infect and reproduce on tomato plants. For this, egg masses were extracted from root-knot galls and examined microscopically to confirm *Meloidogyne* spp. An egg suspension was prepared, and the concentration was adjusted to 2,000 eggs per plant by counting eggs under a microscope using a gridded Petri dish. Now, healthy tomato seedlings were inoculated with eggs into the soil near the root system by making tiny holes around the root zone. After 45 days, the pathogenicity was confirmed through root galling, egg-mass formation, nematode reproduction, and plant symptoms.

### Pathogenicity test of *Fusarium oxysporum*

A pathogenicity test was also conducted to verify the pathogenic nature of *Fusarium oxysporum* culture on tomato plants. A pure culture of the fungus was grown on Potato Dextrose agar (PDA), and a spore suspension was prepared from actively growing cultures. The suspension was standardized to a concentration of 10□ spores mL□¹ using a Neubauer improved hemocytometer. Now, 25 mL of the suspension was applied close to each plant’s root zone. The appearance of characteristic wilt symptoms in inoculated plants, together with the absence of symptoms in the healthy control, was considered evidence of pathogenicity.

### Statistical analysis

Statistical analysis was carried out using a one-factor completely randomized design using R version 4.4.0 in RStudio version 2026.04.0 (Posit, 2026). Analysis of variance (ANOVA) was performed to determine the significance of treatment effects, and treatment means were compared using Duncan’s Multiple Range Test (DMRT) at the 5% significance level using "agricolae" packages. Data processing and visualization were performed using the dplyr, tidyr, and ggplot2 packages. Spearman’s rank correlation analysis and correlation heatmap visualization were performed using the "corrplot" package in R.

## Results

### Vegetative growth

Vegetative growth responses varied among treatments and became more pronounced as the plant aged (Tables 1-3). At 45 DAT, plant height and leaf weight did not differ, whereas shoot, root, and total vegetative weights differed significantly. Plants treated with *T. viride* + *P. fluorescens* produced the highest root weight (6.88 gm) and vegetative biomass (59.63 gm). At 60 DAT, significant differences occurred in plant height, shoot weight, leaf weight, and vegetative biomass, with plants grafted with Solmel being the tallest (116.33 cm) and plants treated with nematode-only recording the highest leaf weight (40.34 gm) and vegetative biomass (112.09 gm). By 75 DAT, all vegetative growth traits differed significantly, with plants treated with *T. viride* + *P. fluorescens* showing high overall biomass, while disease-free plants recorded the greatest plant height (128.67 cm) and root weight (41.60 gm).

**Table 1.** Effect of different bio-management methods on vegetative growth of tomato plants at 45 DAT at Pokhara Kaski, 2026.

| Treatment | Plant height | Stem Weight | Root Weight | Leaf Weight | Vegetative Weight |
| --- | --- | --- | --- | --- | --- |
| *Trichoderma viride* | 49.53 | 28.92 (3.35)<sup>a-c</sup> | 4.46 (1.66)<sup>a-c</sup> | 19.03(1.30) | 47.95(3.83)<sup>ab</sup> |
| *Pseudomonas fluorescens* | 51.53 | 29.40(3.40)<sup>a-c</sup> | 6.15(1.96)<sup>ab</sup> | 16.87(1.25) | 46.27(3.85)<sup>ab</sup> |
| *Trichoderma* + |  |  |  |  |  |
| *Pseudomonas* | 57.20 | 39.11(3.65)<sup>ab</sup> | 6.88(2.04)<sup>a</sup> | 20.53(1.33) | 59.63(4.04)<sup>a</sup> |
| Self-grafted | 44.30 | 17.72(2.90)<sup>b-d</sup> | 2.83(1.29)<sup>c</sup> | 11.35(1.09) | 29.07(3.35)<sup>a-c</sup> |
| *S. sisymbriifolium* | 42.20 | 18.34 (2.91)<sup>b-d</sup> | 3.22(1.44)<sup>bc</sup> | 11.93(1.11) | 32.55(3.50)<sup>a-c</sup> |
| *Solmel* | 45.53 | 19.91(3.02)<sup>b-d</sup> | 2.48(1.20)<sup>c</sup> | 10.20(1.05) | 31.83(3.43)<sup>a-c</sup> |
| *S. torvum* | 40.63 | 14.76(2.71)<sup>cd</sup> | 2.29(1.14)<sup>c</sup> | 15.72(1.22) | 24.96(3.15)<sup>bc</sup> |
| VNRs-10 | 44.60 | 19.18(2.97)<sup>b-d</sup> | 2.50(1.23)<sup>c</sup> | 15.72(1.22) | 34.90(3.53)<sup>a-c</sup> |
| Nematode only (Control) | 48.60 | 31.81(3.47)<sup>a-c</sup> | 6.74(2.03)<sup>a</sup> | 19.51(1.31) | 51.32(3.94)<sup>ab</sup> |
| *Fusarium* only (Control) | 57.00 | 32.01(3.38)<sup>a-c</sup> | 3.70(1.53)<sup>a-c</sup> | 6.39(0.87) | 38.40(3.60)<sup>a-c</sup> |
| Nematode + *Fusarium*<br>only | 51.63 | 14.25(2.48)<sup>d</sup> | 2.83(1.27)<sup>c</sup> | 6.07(0.85) | 20.32(2.76)<sup>c</sup> |
| Disease-free Control | 58.67 | 44.17(3.80)<sup>a</sup> | 2.93(1.35)<sup>c</sup> | 10.58(1.06) | 54.75(4.02)<sup>ab</sup> |
| CV % | 11.82 | 12.75 | 7.61 | 13.75 | 12.50 |
| LSD | - | 0.682 | 0.503 | — | 0.75 |
| F-test | ns | * | ** | ns | . |
| Grand Mean | 49.29 | 25.80 | 3.92 | 13.66 | 39.33 |

**Note:** CV: Coefficient of Variation; LSD: Least Significant Difference; CV% and LSD are of transformed data; Grand mean is calculated from back transformed data; All mean values with same letters (s) are not significantly different; DMRT(p<0.05), ( ): Parenthesis represent transformed data; \*\*\*: significance at 0.1%; \*\*: significance at 1%; \*: significance at 5%;. : significance at 10%; ns: non- significant.

**Table 2.** Effect of different bio-management methods on vegetative growth of tomato plants at 60 DAT at Pokhara Kaski, 2026.

| Treatment | Plant height | Stem Weight | Root Weight | Leaf Weight | Vegetative Weight |
| --- | --- | --- | --- | --- | --- |
| *Trichoderma viride* | 92.83<sup>b</sup> | 58.18(4.08)<sup>a-c</sup> | 12.40(1.13) | 34.71(3.57)<sup>ab</sup> | 92.89 (4.54)<sup>a-c</sup> |
| *Pseudomonas fluorescens* | 84.83<sup>b</sup> | 41.19(3.73)<sup>c</sup> | 11.00(1.08) | 20.66(3.07)<sup>ab</sup> | 61.85(4.13)<sup>b-d</sup> |
| *Trichoderma*+ *Pseudomonas* | 83.67<sup>b</sup> | 50.39(3.93)<sup>a-c</sup> | 10.47(1.06) | 26.00(3.25)<sup>ab</sup> | 76.39(4.34)<sup>a-d</sup> |
| Self-grafted | 95.17<sup>b</sup> | 64.88(4.12)<sup>a-c</sup> | 10.28(1.05) | 24.84(3.00)<sup>a-c</sup> | 89.73(4.40)<sup>a-d</sup> |
| *S. sisymbriifolium* | 96.50<sup>b</sup> | 41.91(3.71)<sup>c</sup> | 14.30(1.18) | 19.23(2.90)<sup>a-c</sup> | 61.14(4.07)<sup>cd</sup> |
| *Solmel* | 116.33<sup>a</sup> | 64.49(4.16)<sup>a-c</sup> | 10.64(1.07) | 26.20(3.30)<sup>ab</sup> | 90.69(4.51)<sup>a-d</sup> |
| *S. torvum* | 98.83<sup>b</sup> | 85.28(4.39)<sup>a</sup> | 11.14(1.08) | 19.97(2.97)<sup>a-c</sup> | 105.26(4.60)<sup>ab</sup> |
| VNRs-10 | 94.83<sup>b</sup> | 53.54(3.98)<sup>a-c</sup> | 11.81(1.11) | 17.29(2.90)<sup>a-c</sup> | 70.83(4.26)<sup>a-d</sup> |
| Nematode only (Control) | 99.33<sup>b</sup> | 71.74(4.28)<sup>ab</sup> | 11.46(1.11) | 40.34(3.70)<sup>a</sup> | 112.09(4.72)<sup>a</sup> |
| *Fusarium* only (Control) | 87.33<sup>b</sup> | 45.11(3.83)<sup>bc</sup> | 10.65(1.07) | 9.05(2.23)<sup>c</sup> | 54.15(4.01)<sup>d</sup> |
| Nematode+ *Fusarium* only | 88.67<sup>b</sup> | 52.06(3.96)<sup>a-c</sup> | 9.51(1.02) | 26.62(3.20)<sup>ab</sup> | 78.67(4.37)<sup>a-d</sup> |
| Disease-free Control | 96.00<sup>b</sup> | 74.84(4.33)<sup>a</sup> | 19.09(1.30) | 14.99(2.77)<sup>bc</sup> | 89.83(4.51)<sup>a-d</sup> |
| CV % | 9.64 | 6.06 | 7.94 | 14.24 | 6.19 |
| LSD | 15.35 | 0.41 | — | 0.74 | 0.46 |
| F-test | * | * | ns | * | . |
| Grand Mean | 94.53 | 58.18 | 11.90 | 23.33 | 81.96 |

**Note:** CV: Coefficient of Variation; LSD: Least Significant Difference; CV% and LSD are of transformed data; Grand mean is calculated from back transformed data; All mean values with same letters (s) are not significantly different; DMRT(p<0.05), ( ): Parenthesis represent transformed data; \*\*\*: significance at 0.1%; \*\*: significance at 1%; \*: significance at 5%;. : significance at 10%; ns: non- significant.

**Table 3.** Effect of different bio-management methods on vegetative growth of tomato plants at 75 DAT at Pokhara Kaski, 2026.

| Treatment | Plant height | Stem Weight | Root Weight | Leaf Weight | Vegetative Weight |
| --- | --- | --- | --- | --- | --- |
| *Trichoderma viride* | 113.83(4.73)<sup>ab</sup> | 60.7 (4.12)<sup>ab</sup> | 20.94(3.04)<sup>b-d</sup> | 14.42(2.72)<sup>b-d</sup> | 75.12<sup>cd</sup> |
| *Pseudomonas fluorescens* | 108.67(4.69)<sup>ab</sup> | 53.19(3.96)<sup>b</sup> | 21.04(2.94)<sup>b-d</sup> | 15.71(2.73)<sup>b-d</sup> | 68.9<sup>cd</sup> |
| *Trichoderma*+<br>*Pseudomonas* | 127(4.84)<sup>a</sup> | 82.95(4.42)<sup>a</sup> | 34.07(3.50)<sup>ab</sup> | 36.54(3.60)<sup>a</sup> | 119.49<sup>a</sup> |
| Self-grafted | 124.33(4.82)<sup>a</sup> | 71.04(4.25)<sup>ab</sup> | 23.81(3.20)<sup>a-d</sup> | 23.53(3.15)<sup>ab</sup> | 94.57<sup>a-c</sup> |
| *S. sisymbriifolium* | 96.67(4.57)<sup>b</sup> | 61.11(4.10)<sup>ab</sup> | 32.27(3.48)<sup>a-c</sup> | 9.28(2.33)<sup>cd</sup> | 70.39<sup>cd</sup> |
| *Solmel* | 115(4.73)<sup>ab</sup> | 76.09(4.34)<sup>ab</sup> | 19.61(3.01)<sup>b-d</sup> | 23.88(3.20)<sup>ab</sup> | 99.97<sup>a-c</sup> |
| *S. torvum* | 117.67(4.76)<sup>a</sup> | 92.57(4.53)<sup>a</sup> | 20.79(3.04)<sup>b-d</sup> | 19.59(3.01)<sup>ab</sup> | 112.17<sup>ab</sup> |
| VNRRs-10 | 110.33(4.70)<sup>ab</sup> | 70.38(4.25)<sup>ab</sup> | 16.78(2.85)<sup>cd</sup> | 19.91(3.03)<sup>ab</sup> | 90.28<sup>a-c</sup> |
| Nematode only (Control) | 118.33(4.77)<sup>a</sup> | 71.74(4.28)<sup>ab</sup> | 22.53(3.15)<sup>a-d</sup> | 24.28(3.23)<sup>ab</sup> | 96.02<sup>a-c</sup> |
| *Fusarium* only (Control) | 96.67(4.57)<sup>b</sup> | 35.68(3.56)<sup>c</sup> | 31.27(3.47)<sup>a-c</sup> | 10.64(2.37)<sup>cd</sup> | 46.32<sup>d</sup> |
| Nematode+ *Fusarium*<br>only | 107.33(4.67)<sup>ab</sup> | 60.39(4.09)<sup>ab</sup> | 12.66(2.60)<sup>d</sup> | 17.78(2.85)<sup>bc</sup> | 78.17<sup>b-d</sup> |
| Disease-free Control | 128.67(4.86)<sup>a</sup> | 70.84(4.27)<sup>ab</sup> | 41.6(3.75)<sup>a</sup> | 7.63(2.13)<sup>d</sup> | 78.47<sup>b-d</sup> |
| CV % | 2.09 | 5.48 | 10.27 | 11.71 | 22.66 |
| LSD | 0.167 | 0.38 | 0.548 | 0.565 | 31.15 |
| F-test | * | ** | * | *** | ** |
| Grand Mean | 113.71 | 67.22 | 24.78 | 18.6 | 85.82 |

**Note:** CV: Coefficient of Variation; LSD: Least Significant Difference; CV% and LSD are of transformed data; Grand mean is calculated from back transformed data; All mean values with same letters (s) are not significantly different; DMRT (p<0.05), (): Parenthesis represent transformed data; \*\*\*: significance at 0.1%; \*\*: significance at 1%; \*: significance at 5%; .: significance at 10%; ns: non- significant.

### Nematode infestation

Nematode infestation differed significantly among treatments at both 60 and 75 DAT (P < 0.001; Tables 4 and 5). Plants grafted with resistant rootstocks consistently showed the lowest galling and nematode reproduction, particularly *S. torvum* and Solmel. At 60 DAT, plants treated with *T. viride* reduced nematode reproduction (RF 8.47; 1,369 eggs gm□¹ root), plants treated with *Pseudomonas fluorescens* produced fewer galls (131.67) but a higher reproductive factor (15.03), whereas the plants treated with *T. viride* + *P. fluorescens* combination showed the highest infestation (320 galls; RF 38.03; 6,951 eggs gm□¹ root). At 75 DAT, plants treated with *S. torvum*, Solmel, and VNRRs-10 grafting maintained very low reproductive factors (0.33, 0.45, and 0.62, respectively), while the plants inoculated with the nematode-only control recorded the highest gall number (555), reproductive factor (126.87), and egg density (11,250 eggs gm□¹ root).

**Table 4.** Effect of different bio-management methods on nematode infestation parameters at 60 DAT at Pokhara, Kaski, 2026.

| Treatment | Gall Number | Gall Index | Reproduction Factor | Egg per gram |
| --- | --- | --- | --- | --- |
| *Trichoderma viride* | 208.00 (14.14)<sup>ab</sup> | 2.67 (1.77)<sup>bc</sup> | 8.47 (3.00)<sup>bc</sup> | 1369.00 (36.99)<sup>b</sup> |
| *Pseudomonas fluorescens* | 131.67 (11.09)<sup>b</sup> | 4.00 (2.06)<sup>ab</sup> | 15.03 (3.20)<sup>bc</sup> | 1867.67 (38.07)<sup>b</sup> |
| *Trichoderma*+ *Pseudomonas* | 320.00 (17.58)<sup>a</sup> | 7.67 (2.86)<sup>a</sup> | 38.03 (6.05)<sup>a</sup> | 6951.00 (82.96)<sup>a</sup> |
| Self-grafted | 20.00 (4.52)<sup>c</sup> | 3.00 (1.84)<sup>bc</sup> | 1.47 (1.39)<sup>cd</sup> | 297.67 (16.90)<sup>bc</sup> |
| *S. sisymbriifolium* | 12.00 (2.48)<sup>c</sup> | 1.67 (1.25)<sup>b-d</sup> | 1.46 (1.20)<sup>cd</sup> | 405.00(12.09)<sup>c</sup> |
| *Solmel* | 1.67 (1.38)<sup>c</sup> | 0.67 (1.05)<sup>cd</sup> | 0.00 (0.71)<sup>d</sup> | 0.00 (0.71)<sup>c</sup> |
| *S. torvum* | 0.00 (0.70)<sup>c</sup> | 0.00 (0.71)<sup>d</sup> | 0.00 (0.71)<sup>d</sup> | 0.00 (0.71)<sup>c</sup> |
| VNRRs-10 | 17.33 (2.89)<sup>c</sup> | 1.33 (1.18)<sup>cd</sup> | 2.23 (1.36)<sup>cd</sup> | 194.33 (8.5)<sup>c</sup> |
| Nematode only (Control) | 209.00 (14.45)<sup>ab</sup> | 7.67 (2.84)<sup>a</sup> | 26.50 (4.98)<sup>ab</sup> | 4218.33 (62.66)<sup>a</sup> |
| *Fusarium* only (Control) | 0.00 (0.71)<sup>c</sup> | 0.00 (0.71)<sup>d</sup> | 0.00 (0.71)<sup>d</sup> | 0.00 (0.71)<sup>c</sup> |
| Nematode+ *Fusarium* only | 238.67 (15.07)<sup>ab</sup> | 7.33 (2.78)<sup>a</sup> | 24.13 (4.84)<sup>ab</sup> | 4935.67 (68.89)<sup>a</sup> |
| Disease-free Control | 0.00 (0.71)<sup>c</sup> | 0.00 (0.71)<sup>d</sup> | 0.00 (0.71)<sup>d</sup> | 0.00 (0.71)<sup>c</sup> |
| CV% | 37.67 | 28.15 | 50.84 | 47.55 |
| LSD | 4.54 | 0.78 | 2.06 | 22.03 |
| F-test | *** | *** | *** | *** |
| Grand Mean | 96.53 | 3 | 9.78 | 1686.56 |

**Note:** CV: Coefficient of Variation; LSD: Least Significant Difference; CV% and LSD are of transformed data; Grand mean is calculated from back transformed data; All mean values with same letters (s) are not significantly different; DMRT(p<0.05), (): Parenthesis represent transformed data; \*\*\*: significance at 0.1%; \*\*: significance at 1%; \*: significance at 5%;.: significance at 10%; ns: non- significant.

**Table 5.** Effect of different bio-management methods on nematode infestation parameters at 75 DAT at Pokhara, Kaski, 2026.

| Treatment | Gall Number | Gall Index | Reproduction Factor | Egg per gram |
| --- | --- | --- | --- | --- |
| *Trichoderma viride* | 76.00 (8.60)<sup>c</sup> | 2.00 (1.56)<sup>cd</sup> | 2.58 (1.71)<sup>cd</sup> | 244.00 (15.34)<sup>c-e</sup> |
| *Pseudomonas fluorescens* | 107.00 (10.08)<sup>c</sup> | 4.67 (2.20)<sup>bc</sup> | 10.35 (2.84)<sup>bc</sup> | 705(24.45)<sup>c</sup> |
| *Trichoderma*+ *Pseudomonas* | 120.00(10.97)<sup>bc</sup> | 7.00 (2.73)<sup>ab</sup> | 8.55 (2.80)<sup>bc</sup> | 471.00 (20.78)<sup>cd</sup> |
| Self-grafted | 12.00 (3.26)<sup>d</sup> | 2.00 (1.56)<sup>cd</sup> | 3.92 (1.86)<sup>b-d</sup> | 340.33(14.40)<sup>c-e</sup> |
| *S. sisymbriifolium* | 23.00 (3.25)<sup>d</sup> | 1.33 (1.17)<sup>de</sup> | 3.67 (1.60)<sup>cd</sup> | 248.33(9.57)<sup>c-e</sup> |
| *Solmel* | 2.67 (1.44)<sup>d</sup> | 0.67 (0.99)<sup>de</sup> | 0.45 (0.92)<sup>d</sup> | 36.67(3.97)<sup>de</sup> |
| *S. torvum* | 3.00 (1.70)<sup>d</sup> | 0.67(1.05)<sup>de</sup> | 0.33 (0.88)<sup>d</sup> | 27.33(3.50)<sup>de</sup> |
| VNRRs-10 | 10.00 (3.11)<sup>d</sup> | 1.00 (1.22)<sup>de</sup> | 0.62 1.04)<sup>cd</sup> | 73.33 (8.37)<sup>c-e</sup> |
| Nematode only (Control) | 555.00 (23.51)<sup>a</sup> | 8.67 (3.02)<sup>a</sup> | 126.87 (11.27)<sup>a</sup> | 11250(106.05)<sup>a</sup> |
| *Fusarium* only (Control) | 0.00 (0.71)<sup>d</sup> | 0.00 (0.71)<sup>e</sup> | 0.00 (0.71)<sup>d</sup> | 0.00 (0.71)<sup>e</sup> |
| Nematode+ *Fusarium* only | 216.00 (14.34)<sup>b</sup> | 7.67 (2.84)<sup>ab</sup> | 13.12 (3.61)<sup>b</sup> | 2320.00 (46.12)<sup>b</sup> |
| Disease-free Control | 0.00 (0.71)<sup>d</sup> | 0.00 (0.71)<sup>e</sup> | 0.00 (0.71)<sup>d</sup> | 0.00 (0.71)<sup>e</sup> |
| CV% | 32.76 | 23.37 | 39.18 | 43.71 |
| LSD | 3.75 | 0.65 | 1.65 | 15.59 |
| F-test | *** | *** | *** | *** |
| Grand Mean | 93.72 | 2.97 | 14.20 | 1309.67 |

**Note:** CV: Coefficient of Variation; LSD: Least Significant Difference; CV% and LSD are of transformed data; The grand mean is calculated from back-transformed data. All mean values with the same letters (s) are not significantly different; DMRT (p<0.05), (): Parenthesis represents transformed data; \*\*\*: significance at 0.1%; \*\*: significance at 1%; \*: significance at 5%; .: significance at 10%; ns: non-significant

### Disease development and management efficiency

Disease development differed substantially among bio-management treatments for both root-knot nematode and *Fusarium* wilt (Figures 1-8). Plants treated with nematode-only and nematode + *Fusarium* recorded the highest nematode AUDPC (235.0 and 232.5) and maintained high disease severity, reaching 73–87% by 45 DAT, whereas plants treated with rootstocks showed consistently low severity and AUDPC. Similarly, *Fusarium* AUDPC was highest under plants treated with nematode + *Fusarium* and *Fusarium*-only (145.0 and 132.5), with severity reaching 93% under combined inoculation. Rootstock treatments maintained the lowest *Fusarium* severity and AUDPC throughout the assessment period. Management efficiency further confirmed the superior performance of rootstocks, with plants grafted with *S. torvum*, Solmel, VNRRs-10, and *S. sisymbriifolium* achieving >85% efficiency against nematodes, while plants grafted with *S. torvum* and VNRRs-10 achieved 92.5% against *Fusarium*. The plants treated with *T. viride* + *P. fluorescens* combination showed limited efficiency against nematodes (16.0%) but moderate efficiency against *Fusarium* (52.8%).

**Figure 1.**
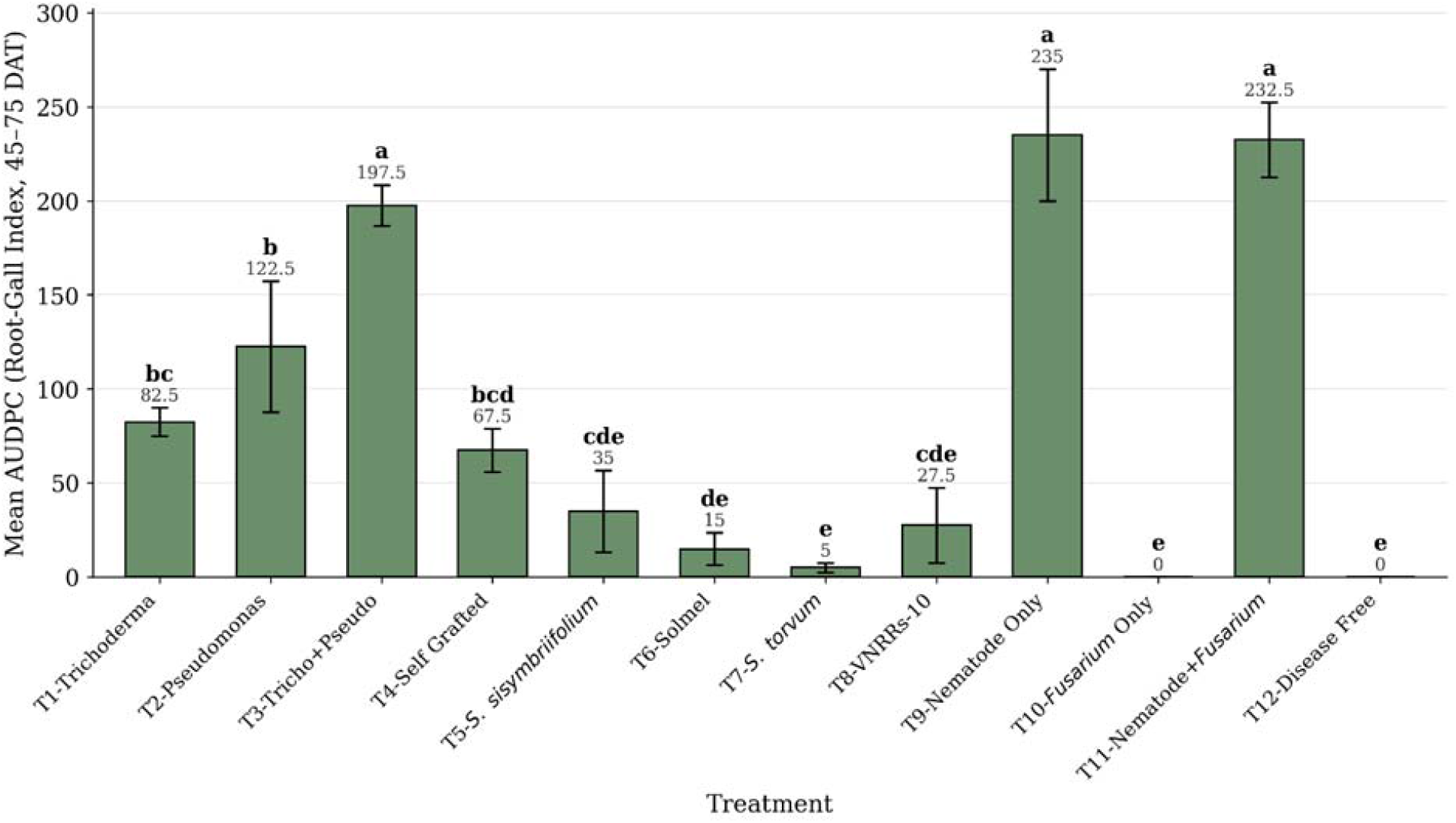
AUDPC of Root-knot nematode severity (Gall index) in Nematode + Fusarium complex under different bio-management methods at Pokhara, Kaski, 2026.

**Figure 2.**
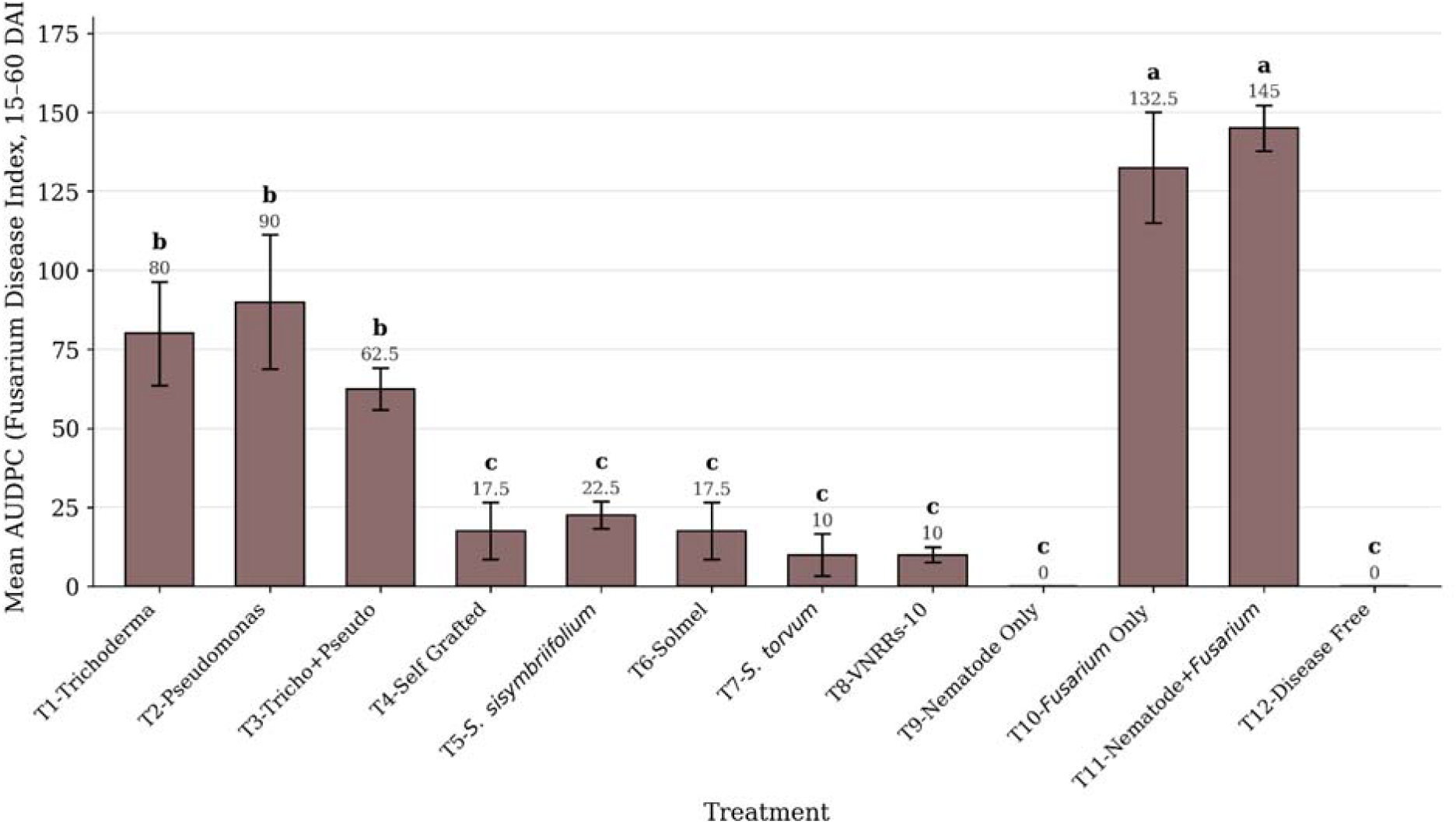
AUDPC of *Fusarium* wilt severity in Nematode + *Fusarium* Complex under different bio-management methods at Pokhara, Kaski, 2026.

**Figure 3.**
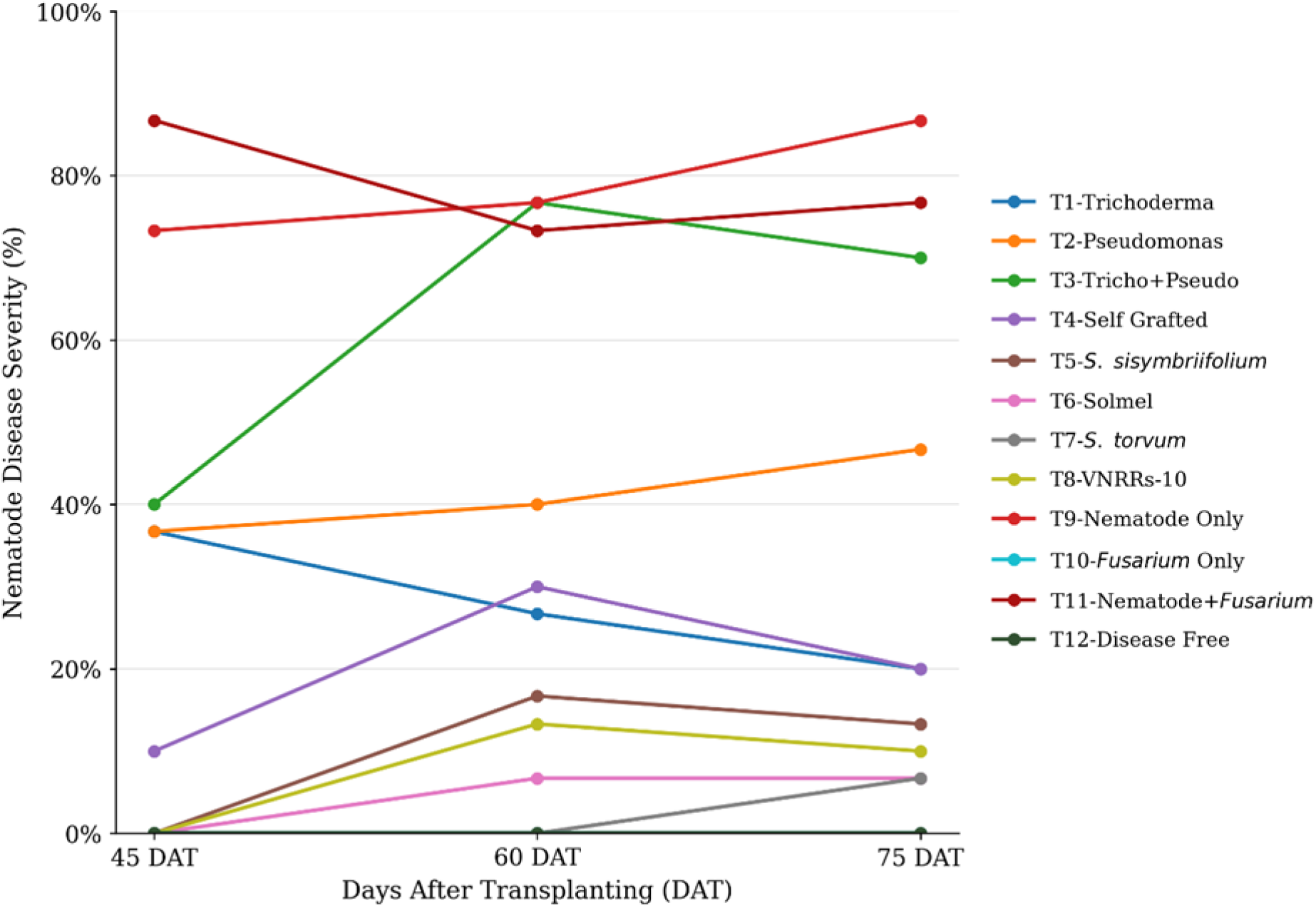
Disease progress curve of Root-knot nematode severity (%) in Nematode + *Fusarium* complex under different bio-management methods at Pokhara, Kaski, 2026.

**Figure 4.**
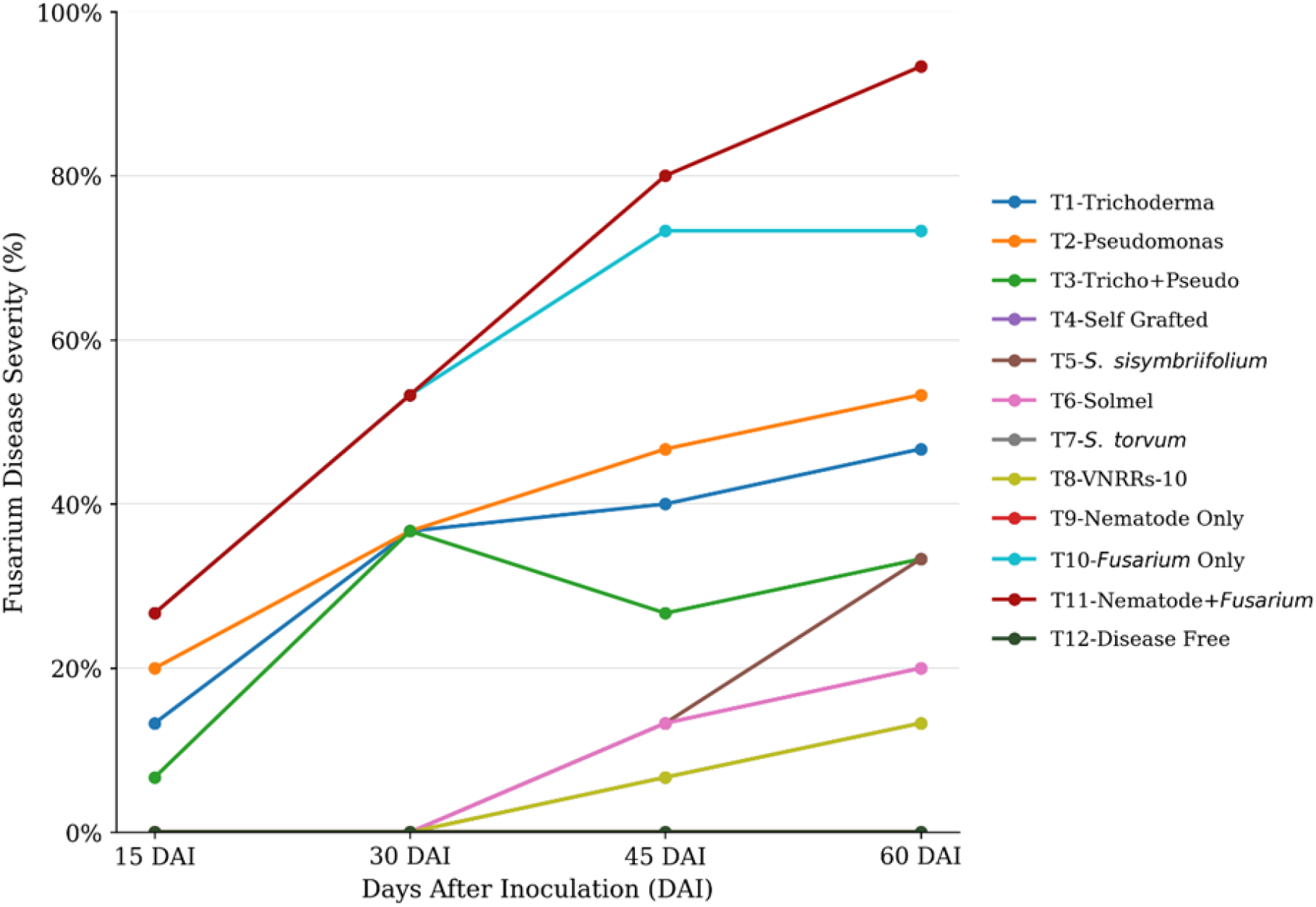
Disease progress curve of *Fusarium* wilt severity (%) in Nematode + *Fusarium* complex under different bio-management methods at Pokhara, Kaski, 2026.

**Figure 5.**
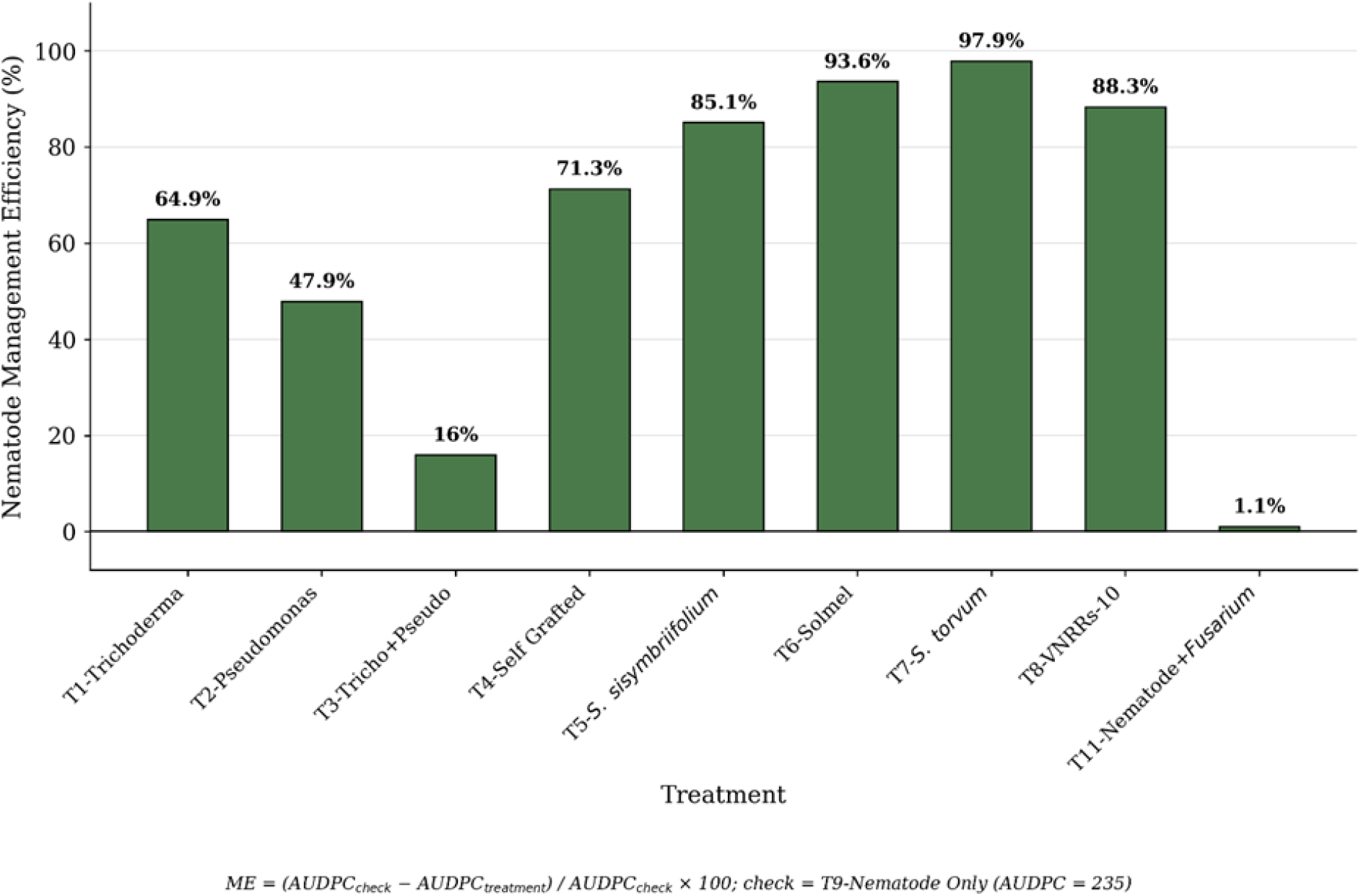
Management Efficiency (%) of different bio-management methods against root-knot nematode infestation in Nematode + *Fusarium* complex at Pokhara, Kaski, 2026.

**Figure 6.**
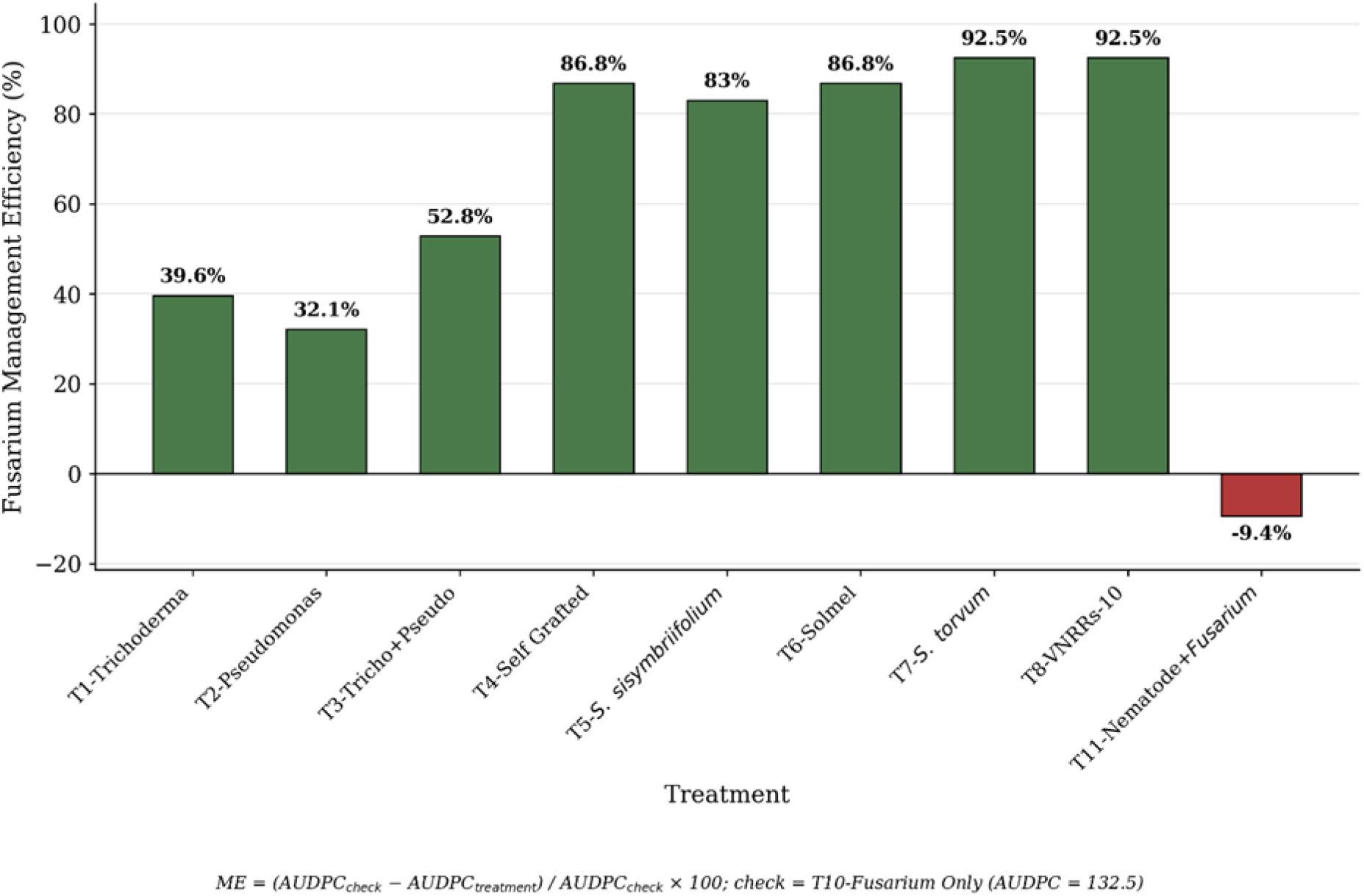
Management Efficiency (%) of different bio-management methods against *Fusarium*infestation in Nematode-*Fusarium* complex at Pokhara, Kaski. 2026.

### Correlation

The correlation between root weight and nematode disease severity is a sharp one-time shift rather than a gradual decline. Root weight correlated positively with all four disease measures (reproduction factor: r = 0.71; gall number: r = 0.62; gall index: r = 0.48; eggs/gram: r = 0.59; all p < 0.01) at 45 DAT, which shifted to non-significant or weakly negative values for every measure by 60 DAT, followed by no shift at 75 DAT (Figure 9-11). The three tests (Fisher’s r-to-z transformation, a permutation test, and an interaction-term regression) confirmed that the change in correlation coefficient from 45DAT to 60DAT is a real shift rather than the statistical illusion caused by the cut-off, whereas no change was noticed for any disease measure. As sampling was destructive and cross-sectional, results showed the shift in population level was within groups instead of within the same plant population.

Plant height validates the weaker version of the same pattern. At 45 DAT, a positive correlation with disease severity (r = 0.15 to 0.22 across the four disease measures), which shifted to negative by 60 DAT. Out of the four measures, three showed significance in this. By 75DAT, the shift did not persist, unlike root weight. The correlation of stem weight and vegetative weight with disease severity stayed the same from 45DAT to 60 DAT to 75 DAT.

Leaf weight in relation to gall number showed a stable, moderate correlation and no significant differences between any pair of coefficients across all three points (r = 0.36 to 0.43; p < 0.05 to 0.01). The asymmetry between root and leaf weight matches the pattern seen in a damage threshold study, which confirmed that as the number of nematodes increases, the root diameter dropped to the minimum lowest value (m = 0.509), while shoot fresh weight recorded the highest yield value (m = 0.761). This supports the idea that root traits were more sensitive to nematode pressure than the shoot traits (Bouchtaoui et al., 2025). It hasn’t been confirmed if this applies to root weight or to eggplant cultivar screening, so it should be taken as a general comparison rather than solid proof.

**Figure 7.**
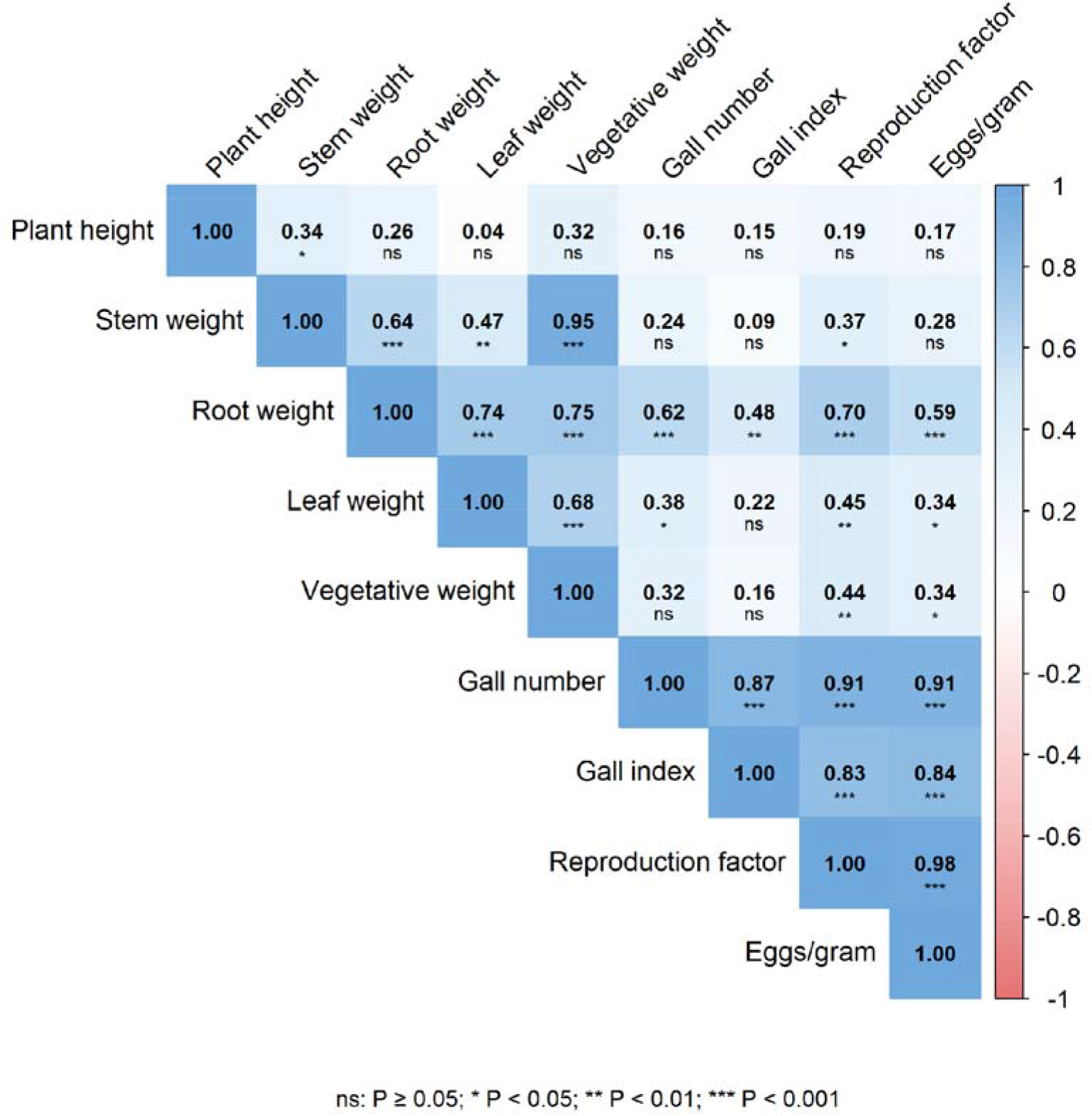
Spearman correlation heatmap showing the relationships between vegetative growth traits and root-knot nematode infestation parameters at 45 DAT at Pokhara, Kaski. 2026.

**Figure 8.**
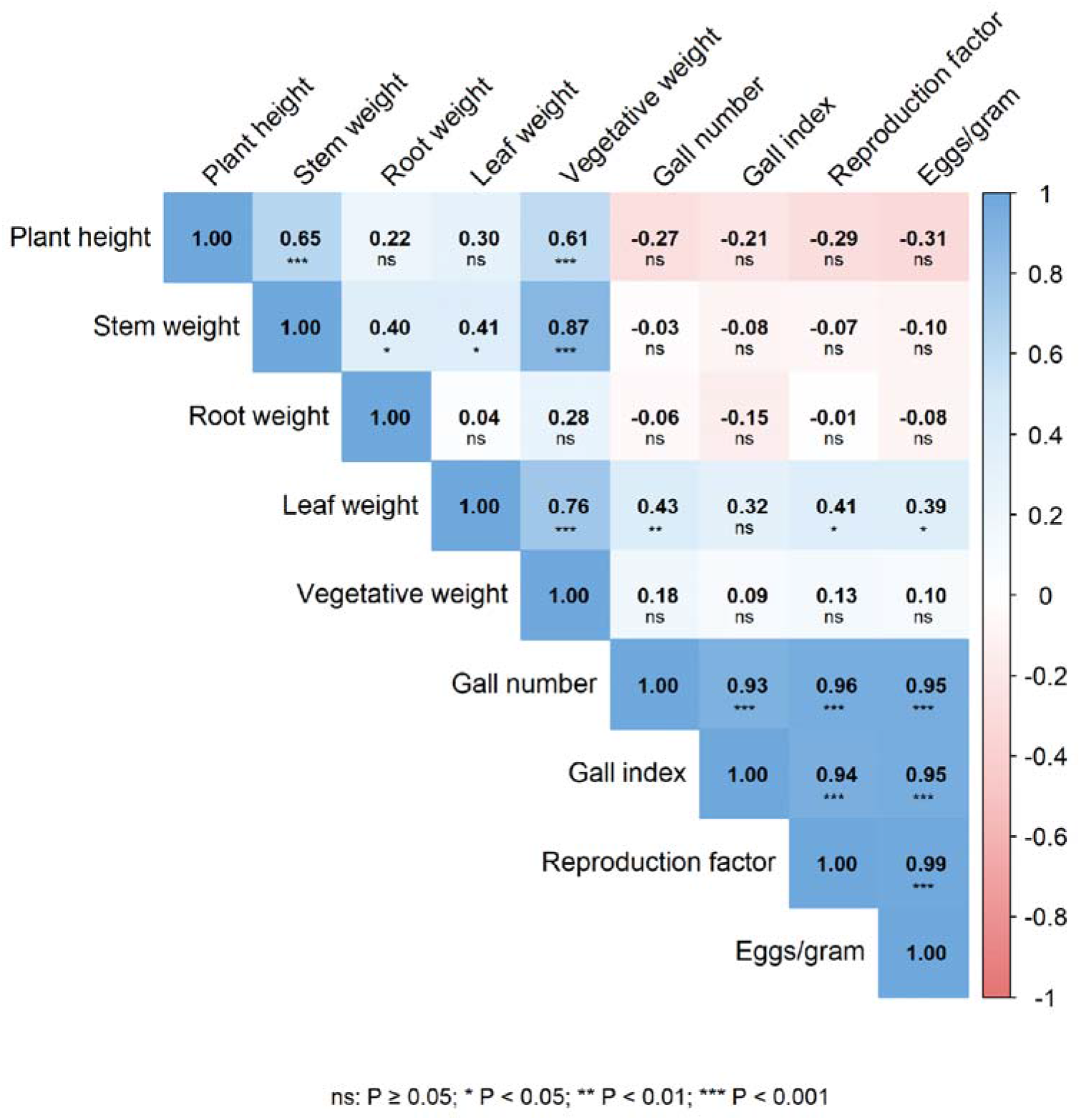
Spearman correlation heatmap showing the relationships between vegetative growth traits and root-knot nematode infestation parameters at 60 DAT at Pokhara, Kaski. 2026.

**Figure 9.**
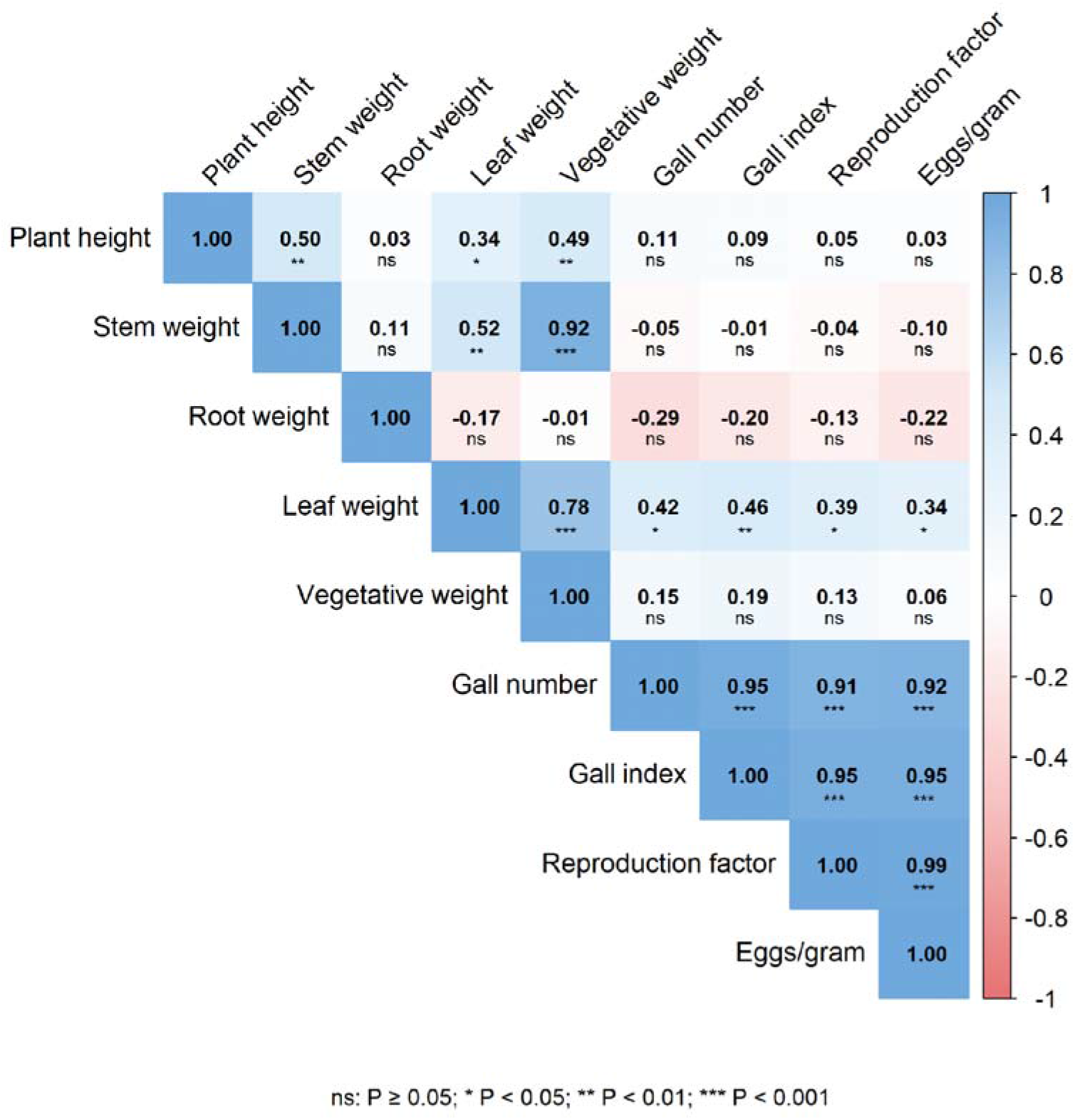
Spearman correlation heatmap showing the relationships between vegetative growth traits and root-knot nematode infestation parameters at 75 DAT at Pokhara, Kaski. 2026.

## Discussion

### Vegetative growth response

#### Effect of rootstocks

Across all three sampling dates the resistant rootstocks effectively suppressed nematode infestation; however, the resistance that they carried was not consistently accompanied by the larger gains in the vegetative growth. At 45 DAT, several rootstocks even recorded comparatively low root and shoot biomass, suggesting that the resistance of the rootstocks has no positive association with their early plant vigor. The obtained observation outcome is not a contradiction, but it likely reflects the transient growth lag that is associated with graft union establishment (Nordey et al., 2020), which delays growth by roughly one to three weeks, resulting in a deficit in cumulative growing time. physiological cost of establishing graft union. Even among broadly compatible species, there may initially be temporary vascular discontinuity at the graft union, restricting the translocation of water and nutrients until functional vascular reconnection is established between them (as disconnected xylem is separated by parenchyma tissue that fills in only progressively over the first 10-20 days after the grafting) (Frey *et al*., 2020b). This discontinuity translates into measurable hydraulic resistance at the union itself: in tomato grafted specifically onto *Solanum torvum* rootstock, shoot thickening, xylem distortion, and reduced water potential immediately above the graft interface were all greatest relative to tomato or eggplant rootstocks, restricting water and nutrient translocation to the scion until functional vascular reconnection is established (Oda *et al*., 2005).

By the time of the 3rd sampling (75 DAT), this effect had been substantially recovered, with self-grafted and other rootstock-grafted plants approaching or exceeding the disease-free control for plant height and root weight. An important practical implication that may be associated with this finding is that the resistant rootstock should not be evaluated solely based on its early vegetative performance, as the cumulative benefit of the rootstock may become more apparent over time through suppression of the Nematode-*Fusarium* disease complex. The enhanced resistance in the rootstocks may be associated with structural and physiological defense mechanisms, including lignification, phloem formation, and restricted pathogen movement within the vascular tissue (Beckman, 1987; Rodriguez-Molina, 2003).

### Effect of *Trichoderma viride* and *Pseudomonas fluorescens*

*T. viride* and *P. fluorescens* applied alone produced vegetative performance that tracked closely with the nematode-only control rather than the disease-free control, particularly at 45 and 60 DAT. This is consistent with their principal mode of action being disease suppression and growth promotion via indirect mechanisms - siderophore-mediated iron competition, phosphate solubilization, and phytohormone production (Reddy, 2025) - rather than the constitutive structural resistance conferred by grafting. Growth promotion from these agents depends on successful rhizosphere colonization, which is a probabilistic, environment-sensitive process, and this likely explains why single-agent treatments showed more temporal variability in ranking across the three sampling dates (45, 60, 75 DAT) than the rootstock treatments.

### Combined bio-control treatment

The combined *T. viride* and *P. fluorescens* treatment produced the strongest vegetative growth of any treatment by 75 DAT, exceeding even the disease-free control on leaf and vegetative weight. This is a biologically important and easily overlooked result because it is not consistent with the same treatment being comparatively weak in nematode suppression, discussed in the next part of this discussion section. The two outcomes are not contradictory once growth promotion and pathogen suppression are recognized as least partially separable functions of this organism. Plant growth promotion activity, IAA gibberellin production, improved root architecture, and nutrient acquisition (Reddy, 2025; Vinale *et al*., 2008) do not require the same degree of rhizosphere dominance or sustained colonization density that direct antibiosis or induced resistance against nematode reproduction does. Priming-based induced systemic resistance, one of the principal mechanisms to which both *T. viride* and *P*. *fluorescens* act on the plant immune system, imposes a comparatively low continuous metabolic burden on colonizing organisms related to direct antibiosis or mycoparasitism (Pieterse *et al*., 2014). This may further explain why a growth benefit persisted even when nematode suppression did not. In other words, the combined treatment may have retained enough of each organism’s growth-promoting activity to drive superior biomass accumulation, even while the reduced per-agent inoculum density, limited its capacity to suppress nematode multiplication of the same degree as either agent applied at full strength.

The Nematode + *Fusarium* complex treatment had the lowest vegetative mass at every sampling date (45, 60, 75 DAT), and the magnitude of this reduction was greater than that caused by either of pathogen applied alone. This is the expected consequence of compounding mechanical and physiological injury. Root Knot-nematode feeding induces giant-tissue cells that act as persistent metabolic sinks and physically compromise the root cortex (Eisenback, 2022; Perry *et al*., 2009). While, the resulting wound and altered root exudate profiles due to injury are known to facilitate more rapid fungal ingress and colonization by *Fusarium* (Kawamura & Hirano, 1968; Morrell & Bloom, 1981), Once inside the vascular tissue, *Fusarium* triggers tylose and gum deposition that obstruct the water and nutrient transport along xylem vessels (Agrios, 2005; Beckman, 1987). This vascular restriction is far more consequential for shoot biomass than nematode feeding alone, which explains why *Fusarium*-only control also recorded disproportionately low leaf weight relative to its plant height at 60 DAT. Rootstock that maintained growth under this pressure did so by denying the nematode an initial entry point rather than by tolerating the complex ones established. Resistance expressed at the level of gall initiation removes the wounding step that the fungal component of the complex appears to exploit.

### Nematode infestation

The pattern across all four nematode parameters was consistent. Rootstock suppressed the nematode infestation more completely. The nematode-only control carried the heaviest burden by 75 DAT. The bio-control treatment occupied an intermediate position that shifted between sampling dates. The near-zero gall count on plants grafted with *S. torvum* and Solmel are consistent with the resistance mechanism already documented for this genotype, in which localized hypersensitive response at the site of juvenile penetration prevents giant cell formation before a feeding site can be established. (El-Sappah *et al*., 2019; Wubie & Temesgen, 2019). Because reproduction factor depends entirely on the successful establishment of the feeding site, a rootstock that blocks penetration produces the appearance of suppressing every downstream parameter simultaneously, which is exactly the pattern observed.

The near immunity of *S. torvum* and Solmel relative to the more moderately suppressed resistance seen in the *S. sisymbriifolium* and VNRRs-10, points to the difference in the strength or completeness of the resistant mechanism rather than a difference in the kind. *S. sisymbriifolium* has been reported to confer resistance predominantly through elevated phenolic and defense enzyme accumulation post penetration (Baidya *et al*., 2017), a mechanism that can reduce reproduction without fully preventing initial gall formation, consistent with low but non-zero gall counts recorded on the rootstock in this present trial.

The higher gall number, gall index, and reproduction factor recorded in the combined *T. viride* and *P. fluorescens* treatment than in untreated nematode control at 60 DAT is the strangest result of this study. One of the simplest possible explanations for this can be the application dose, as each bioagent was applied at half the concentration used in the single-agent treatment. Since colonization density drives suppressive capacity for both organisms, the lower inoculum alone could account for weaker performance. However, such pronounced severity also suggests the occurrence of microbial incompatibility between the bioagents. *P. fluorescens* produces antifungal metabolites like 2,4-diacetylphloroglucinol and phenazine-1-carboxylic acid, which possess broad-spectrum activity and are capable enough to affect *Fusarium oxysporum* itself (Raaijmakers, 2004; Rompalli *et al*., 2016). Hence, those compounds could also inhibit *T. viride* establishment when under the same rhizosphere region. Direct evidence of such compatibility incompatibility was provided by Pan & Jash (2010), reporting inhibition of *T. viride* growth by *P. fluorescens* of up to 59%, depending on the bacterial isolate and sequence of inoculation. Similarly, Jambhulkar *et al*. (2018) found synergistic interaction in only one of the six evaluated biocontrol agent combinations, with the remaining combinations exhibiting additive or antagonistic effects. This finding supports the view that the antagonistic interactions among biocontrol agents are more common than that of true synergism. Likewise, a review done by Xu *et al*. (2011) also reported antagonisms are more frequently occurring than that of the synergy. Although some strain-specific combinations of *T. viride* and *P. fluorescens* have shown improved disease suppression (Cai *et al*., 2021), compatibility is highly dependent on the strains that are involved and the external environmental conditions. Therefore, the reduced efficacy of the combined biocontrol treatment is more convincingly attributed to documented antagonistic interactions between *T. viride* and *P. fluorescens* than to an isolated or anomalous response. These findings highlight the importance of evaluating microbial compatibility prior to co-application, as combining individually effective biocontrol agents does not necessarily result in additive or synergistic disease suppression.

### AUDPC and disease progression

#### Nematode AUPDC

Nematode AUDPC mirrors the gall index and reproduction factor results: the combined treatment was statistically indistinguishable from untreated checks across the full period, single agents achieved intermediate suppression, and rootstocks most completely *S. torvum—matched* the pathogen-free condition. Because AUDPC integrates the full assessment window, this indicates the combined treatment’s poor performance was sustained, not a transient effect that resolved later, consistent with the dose-limitation explanation above.

These findings are only partly consistent with previous reports. The intermediate suppression achieved by single biocontrol agents and the near-complete resistance of *S. torvum* agree with the wider literature: Kamalanathan *et al*. (2023) and Poonia *et al*. (2025) similarly report single *Trichoderma* or *Bacillus* isolates producing substantial but incomplete RKN suppression in tomato, and *S. torvum’s* status as a highly resistant rootstock, with nematode multiplication reduced to a small fraction of that on susceptible rootstocks, is well established (Fullana *et al*., 2024; Zhang *et al*., 2023). The combined-treatment result is harder to reconcile with prior work. Studies combining *Trichoderma* sp. and *Pseudomonas* sp. or related multi-agent consortia generally report the combination performing no worse than the better single agent, not collapsing to check-level performance. Mokhtari *et al*. (2014) found *T. harzianum* + *P. fluorescens* CHA0 significantly reduced nematode indices relative to *P. fluorescens* alone and the untreated control, with no significant difference from *T. harzianum* alone. Varkey *et al*. (2018), reviewed in Kumar and Dara (2021), similarly found that combining *Piriformospora indica* with *Bacillus pumilus* and *P. fluorescens* gave no added suppressive benefit over the single agents, and that *Piriformospora indica’s* suppressive effect was actively reduced by antagonism from the bacterial partners, evidence that negative interactions between biocontrol agents, while less commonly reported than synergy, are a recognized outcome. None of these studies, however, report a combined treatment regressing all the way to untreated-check performance. This suggests the antagonism observed here was more severe than the mild negative interactions typically documented in the literature, which reinforces rather than merely supports the dose-limitation explanation offered above: The halved inoculum is likely doing most of the explanatory work for a result this pronounced.

### Fusarium AUPDC within the complex

*Fusarium* AUDPC did not differ significantly between the *Fusarium*-only and Nematode-*Fusarium* complex, which appears to conflict with documented synergism between these pathogens (Göze Özdemir *et al*., 2022; Khan & Sharma, 2020). This is likely because AUDPC represents a season-long cumulative disease development measure over the entire assessment period, whereas nematode-driven acceleration of *Fusarium* wilt infection is greatest during the early-to-mid window when nematode wounds provide an efficient entry route (Morrell & Bloom, 1981; Kawamura & Hirano, 1972). As the disease progressed, both treatments reached similar final disease levels, reducing the impact of early differences on cumulative AUPDC value. In contrast, biocontrol treatments showed intermediate AUDPC values, suggesting that *T. viride* suppressed through direct mycoparasitism, while P. *fluorescens* induced systemic resistance, together providing greater protection against *Fusarium* wilt.

### Disease Severity

#### Fusarium-Nematode interaction and the basis of synergism

The higher disease severity in the Nematode-Fusarium complex compared with *Fusarium-only* treatment at 60 DAT exceeds provides clear evidence for pathogen synergism. This interaction occurs because of the nematode feeding the root tissue, creating a wound that facilitates fungal entry, altering the root tissue in the ways that favor fungal colonization and weaken the plant defense system (Meena *et al*., 2021; Nelson, 1991; Steinkellner *et al*., 2008). As a result, plants infected with both diseases develop more severe disease than those infected with *Fusarium* alone. The findings indicate that early nematode management is essential, as it limits the *Fusarium* infection and reduces the severity of the nematode-Fusarium disease complex.

### Management Efficiency

#### Grafting versus biocontrol

Rootstock consistently performed better than biocontrol across both disease components because the two strategies differ structurally in a way resistance is delivered. The resistance mechanism of rootstock is inherent resistance that is stable throughout the growing season, whereas biocontrol efficacy depends on successful establishment of beneficial microorganisms in the rhizosphere and is influenced by environmental conditions and application dose. This dependence on establishment is the principal reason why biocontrol is more variable in efficacy trials than resistant rootstocks (Ajilogba & Babalola, 2013; Meshram & Adhikari, 2024); however, this variability doesn’t reduce the validity of biocontrol in disease management. Instead, biocontrol functions best as a complement to genetic resistance, an approach increasingly reflected in strategies combining grafting with microbial biocontrol precisely because the two address different points of pathogen vulnerability (Nagachandrabose *et al*., 2025).

### Correlation

The root weight findings noticed could be because of the destructive sampling designs where the group of plants measured at each time were different, and the composition of the groups was also different, rather than a physiological decline within an individual plant. This is the interpretation that was drawn. Second-stage juveniles of *Meloidogyne* establish permanent feeding sites within the root vascular cylinder, which triggers the formation of galls through localized cell division and enlargement (Vilela et al., 2025). The resulting gall tissue mass relative to root biomass has been used as an indicator of symptom severity in root-knot disease (Singh et al., 2006). Since measured whole-root weight instead of gall tissue and functional root weight, and different groups of increasingly diseased plants were measured, this could have led to the weighing of gall swelling rather than the true root. This confound could erode the correlation between root weight and disease severity. So, the mechanism should be read as a plausible interpretation rather than a demonstrated one. This interpretation aligns with the findings of Zhang et al (2006) that gall severity is associated with negative plant growth, and eggplant infected with *Meloidogyne similarly exhibited reduced growth and biomass accumulation* (Vilela et al., 2025).

Very high correlations were observed among gall number, gall index, reproduction factor, and eggs per gram (r = 0.91 to 0.99 across all time points). This reflects the fact that the reproduction factor and eggs/gram are derived from gall and egg counts according to Oostenbrink (1966); Taylor & Sasser (1978). This overlap simply confirms that our measurements are internally consistent rather than representing four independent findings.

These results suggest that once RKN infection is established, root weight is no longer a trustworthy way for detecting plant vigour. This is mainly because, as the disease progresses, more of the weight is contributed by the disease-related swelling rather than the actual growth. Treatments that appear to preserve root weight could be masking the galling rather than reflecting the real tolerance. Leaf weight, on the other hand, could be a more reliable factor for comparing treatments, especially after the early-stage infection, as leaf weight measured a step removed from the infection site isn’t affected by the same problem. While choosing root weight as a measurement, it is advisable to pair root weight with gall index rather than root weight alone.

The principal limitation of his design is that each time point involved a separate group of plants rather than the same plants measured repeatedly. Because of this, the pattern observed mainly describes the changes in the association of a group of plants rather than the disease progression within an individual plant. So, a non-destructive longitudinal design would be needed to test the latter directly. One plant (T11, Rep 1, the RKN + *Fusarium* complex treatment) at 45 DAT could not have its gall number, reproduction factor, and eggs/gram measured; although it had the maximum gall index score, its roots had completely disintegrated. This observation was excluded pairwise (not zero-imputed) from the affected variables, and a sensitivity check confirmed it does not disproportionately drive the reported correlations. No earlier studies have reported this kind of decay in the correlation between root weight and reproduction parameters across sequential destructive harvests. While the literature that we had cited supports the idea that our explanation is reasonable, it does not demonstrate this pattern in itself.

## Conclusion

The study confirmed that the root-knot nematode–*Fusarium oxysporum* disease complex caused greater damage severity to Srijana tomato than either pathogen alone, demonstrating a higher severity effect. Among the resistant rootstocks, *Solanum torvum* was the most effective overall, while *Solanum sisymbriifolium alongside* Solanum torvum provided the greatest suppression of *Fusarium* wilt. Among the biocontrol agents, *Trichoderma viride* performed better than *Pseudomonas fluorescens*, although biological control was less effective than grafting. Therefore, grafting Srijana tomato onto *Solanum torvum* is recommended as the most effective and sustainable management strategy for the root-knot nematode–*Fusarium* disease complex under polyhouse conditions.

## Acknowledgments

The authors would like to express profound gratitude to all the farmers and other stakeholders who provided valuable information for this study.

## Funding

This research was funded by LIBIRD Nepal.

## Conflict of Interest

The authors declare that there is no conflict of interest with regard to this manuscript publication.

## Authors’ contribution

AB was the principal investigator and led the study from the conception of the research idea to the preparation of the final manuscript, including on-the-ground fieldwork and implementation activities, data collection, and analysis. AB (1), AB (2), SG, PS, SN, AB, DK carried out the fieldwork, data collection, data visualization, and formal analysis. SD and SA provided academic guidance and mentorship throughout the entire research and contributed to Manuscript preparation.

## Data availability statement

The data that support the findings of this study are available from the corresponding author upon reasonable request.

## About the Authors

**Anup Bhatt** holds a bachelor’s degree in agricultural science from Agriculture and Forestry University, Rampur, Chitwan, Nepal.

**Aditi Bhandari** holds a bachelor’s degree in agricultural science from Agriculture and Forestry University, Puranchaur, Kaski, Nepal.

**Sajan Gaudel** holds a bachelor’s degree in agricultural science from Agriculture and Forestry University, Rampur, Chitwan, Nepal.

**Ankita Baral** holds a master’s degree in plant pathology from Agriculture and Forestry University, Rampur, Chitwan, Nepal.

**Sparsha Naharki** holds a bachelor’s degree in agricultural science from Agriculture and Forestry University, Rampur, Chitwan, Nepal.

**Priya Shrestha** holds a bachelor’s degree in agricultural science from Agriculture and Forestry University, Rampur, Chitwan, Nepal.

**Deepa Koirala** is currently studying at Agriculture and Forestry University, Rampur, Chitwan, Nepal.

**Suman Dhakal** serves as Assistant Professor at the Department of Agronomy, Faculty of Agriculture, Agriculture and Forestry University, Rampur, Chitwan, Nepal.

**Shalik Ram Adhikari** serves as Chief of the Provincial Plant Protection and Pesticide Management Laboratory, Gandaki Province, Pokhara, Kaski.

## Notes

### Competing Interest Statement

The authors have declared no competing interest.

